# Succession of bacterial communities on carrion is independent of vertebrate scavengers

**DOI:** 10.1101/744748

**Authors:** Cody R. Dangerfield, Ethan Frehner, Evan Buechley, Çağan H. Şekercioğlu, William J. Brazelton

## Abstract

The decomposition of carrion is carried out by a suite of macro- and micro-organisms who interact with each other in a variety of ecological contexts. The ultimate result of carrion decomposition is the recycling of carbon and nutrients from the carrion back into the ecosystem. Exploring these ecological interactions among animals and microbes is a critical aspect of understanding the nutrient cycling of an ecosystem. Here we investigate the potential impacts that vertebrate scavenging may have on the microbial community of carrion. In this study, we placed seven juvenile domestic cow carcasses in the Grassy Mountain region of Utah, USA and collected tissue samples at periodic intervals. Using high-depth environmental sequencing of the 16S rRNA gene and camera trap data, we documented the microbial community shifts associated with decomposition and with vertebrate scavenger visitation. The remarkable scarcity of animals at our study site enabled us to examine natural carrion decomposition in the near absence of animal scavengers. Our results indicate that the microbial communities of carcasses that experienced large amounts of scavenging activity were not significantly different than those carcasses that observed very little scavenging activity. Rather, the microbial community shifts reflected changes in the stage of decomposition similar to other studies documenting the successional changes of carrion microbial communities. Our study suggests that microbial community succession on carrion follows consistent patterns that are largely unaffected by scavenging.

## Introduction

Carrion, or dead animal tissue, provides a nutrient-rich resource for a wide array of organisms. At the smallest scale, both geographically and in organisms affected, carrion contributes nutrients to soils via nutrient leaching, thereby affecting microbial communities in soil near the carcass (Howard, Duos, and Watson-Horzelski 2010; Parkinson et al. 2009; Parmenter and MacMahon 2009). The impacts of carrion can be seen more directly as a food source to the many necrophagous arthropods and vertebrate scavengers (Jordan et al. 2015). Moreover, larger-scale impacts of carrion have been well documented. For example, the massive die-off of salmon and cicada lead to large increases in resources and nutrient availability that affect a myriad of organisms including microbes, plants, fungi, and vertebrates (Hocking and Reynolds 2011, 2012; Jordan et al. 2015; Tiegs et al. 2009, 2011; Yang 2004) The versatile methods by which carrion can be produced and consumed gives it the potential to impact many facets of an ecosystem, but the pathway by which it decomposes and enters the ecosystem’s nutrient cycle depends on the environmental conditions and the interactions that form between the organisms that compete over its resources.

The decomposition of carrion occurs continuously (Schoenly and Reid 1987), but it typically has a consistent progression that is categorized into five stages of decay based on physical composition of the carcass: fresh, bloat, active decay, advanced decay, and putrid dry remains (Payne 1965). A body enters the fresh stage at death when a depletion of internal oxygen triggers autolysis of the cells. Concurrently, endogenous microbes then begin to metabolize the body and produce volatile compounds. The carcass transitions to the bloat stage as the activity of these microbes fill the body cavity with gases, causing the carcass to distend. Active decay follows the bloat stage when the body cavity ruptures, releasing the gases and allowing invertebrates to consume soft tissue within the body cavity. After the removal of most of the soft tissue and decrease in invertebrate activity, the carcass transitions into the advanced decay stage. The carrion enters the putrid dry remains stage when the carcass has desiccated and all that remains are bones and small amounts of skin and hair (Goff 2009; Payne 1965).

The high nutrient content of carrion makes it a highly sought-after resource for many organisms (Hanski 1987; Janzen 1977; Wilson and Wolkovich 2011). Due to the high level of competition, the organisms that consume carrion have developed behaviors in order to monopolize the nutrients of carcass for themselves. These complex interactions among microbes and scavenging fauna, along with abiotic factors (e.g. precipitation and temperature), often impact the duration and occurrence of the stages of decomposition (Carter, Yellowlees, and Tibbett 2010, 2008; Comstock et al. 2015; Galloway, Jones, and Parks 1989; Payne 1965; Rozen, Engelmoer, and Smiseth 2008; Shukla et al. 2017). After an animal’s death, microbes quickly colonize the carcass and begin to metabolize tissue while producing toxins in order to hinder consumption from other organisms (Burkepile et al. 2006; Janzen 1977; Rozen et al. 2008). Vertebrate scavengers, especially vultures, seek to quickly locate and consume carcasses before other scavengers consume them or before decomposition progresses (Buechley and Sekercioglu 2016). Furthermore, some vultures, such as the Turkey Vulture (*Cathartes aura*), have developed an unusually high tolerance to decomposer-produced toxins, such as botulism, and the harsh conditions present in their hindgut reduce the likelihood of carrion microbes surviving consumption and infecting the vulture itself (Beasley, Olson, and DeVault 2015; DeVault, Rhodes Olin E., and Shivik 2003; Roggenbuck et al. 2014). Other behaviors such as the burial of carcasses has developed in both vertebrate and invertebrate scavengers in order to seclude the carrion from climatic conditions, microbes, and other scavengers to slow decomposition and secure the resources for themselves (Frehner et al. 2017; Rozen et al. 2008; Shukla et al. 2017). In addition to burying, Burying Beetles (*Nicrophorus spp.*) further suppress competition with microbes by excreting antimicrobial exudates on the carcass In doing so, these beetles limit decomposition and can delay the carcasses from entering the bloat or active decay stages (Shukla et al. 2017), which are mainly dictated by microbial and insect activity (Finley, Benbow, and Javan 2015; Goff 2009; Payne 1965).

Researching these interactions is important to understand how carrion decomposition impacts nutrient cycling and the importance that carrion has on ecosystems (Barton 2015; Barton et al. 2013), and also to forensics, as the pattern of succession on carrion and cadavers can be used to determine a postmortem time interval (PMI; Anderson 2015). Historically, a majority of studies focused on forensic entomology to determine PMI (Byrd and Allen 2001; Michaud and Moreau 2009; Payne 1965; Schoenly and Reid 1987). Recent studies have utilized DNA sequencing technology to characterize the microbiome of carrion and investigate the potential use of microbes as indicators for PMI (Guo et al. 2016; Hyde et al. 2013; Pechal et al. 2014, 2013). These studies have investigated the microbial communities associated with carrion decomposition and how seasonal changes and macroinvertebrates impact those microbial communities. Moreover, animal scavengers may also impact the microbial composition of the carcass by introducing their own communities of microbes. Scavengers act as a vector of dispersal for many microbes, so their presence or absence may have a significant impact on the microbial community of carrion (Crippen, Benbow, and Pechal 2015).. However, to our knowledge, no study has investigated the influence of scavenger activity on the microbial community composition of carrion. In this study, we use environmental DNA sequencing and vertebrate scavenging data to investigate decomposition dynamics and potential impacts that vertebrate scavengers have on the microbiome of carrion in the Great Basin Desert of Utah.

## Methods

### Study sites and field data collection

In this study, we investigate the bacterial communities of bovine carcasses in the Grassy Mountain region of Utah, USA (40.87°N, -113.03°W) from May to June, 2015. To do so, we experimentally placed juvenile domestic cow (*Bos taurus*) carcasses (n=7) in the study site and monitored their decomposition using camera traps to identify vertebrate scavenger activity and by collecting tissue samples at regular intervals to identify progression of microbial communities. The calves were obtained from one local Utah dairy and had died from natural causes either during or shortly after birth. The carcasses were collected on the day of birth/death, and were kept frozen until their placement in the field to minimize any decomposition progression. They were placed at least 3 km apart and fixed to a concealed stake in the ground to prevent scavengers from removing the complete carcass. The carcasses weighed between 18.6 and 26.9kg. Carcasses were placed on sites that included sparse Utah juniper (*Juniperus osteosperma*), greasewood (*Sarcobatus vermiculatus*), and widely distributed cheatgrass (*Bromus tectorum*). The soil in the study area is composed of loose to moderately compacted limnological sediments, including gravels and clays. The study area is arid and largely homogenous. Study area temperatures varied between 7-40°C, and there was no precipitation during the experiment. We collected tissue samples from each of the carcasses during five sampling periods (Day 1, Day 4, Day 12, Day 18, and Day 26) (Fig 1).

**Figure 1:**
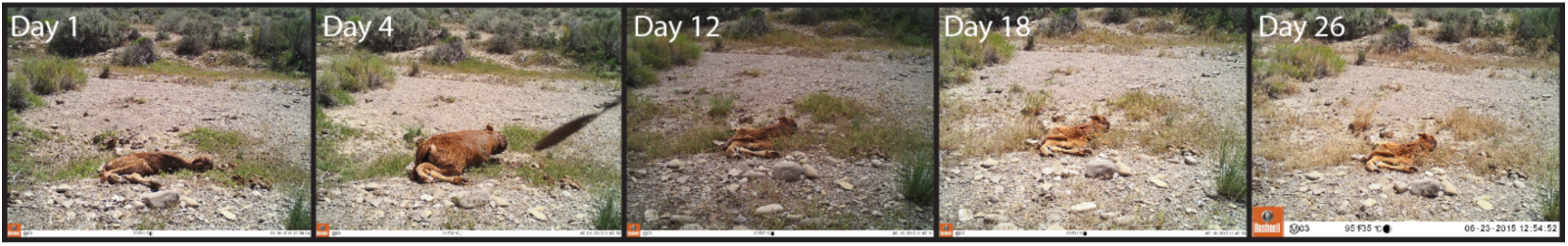
Photographs of the decomposition progression at one of the seven sites. The photos from this site were selected for clarity purposes, and all other sites had similar carcass composition.

The carcasses were equipped with Bushnell Trophy Cam HD motion-activated cameras to monitor vertebrate scavenging activity. The cameras were programmed to take 1 photo when triggered, with a 10-s delay between subsequent photos to reduce saturation of photos from the same animal visitation event. All photos collected over the course of the study were entered into CameraBase Version 1.7 (Tobler 2007), a camera-trap photo management platform in Microsoft Access. We analyzed each of these photos individually and identified any vertebrates present in the photos to species. We identified arrival times after carcass placement and duration of presence at carcass for each scavenger species.

Tissue samples were excised from hind-leg muscle tissue of each carcass. Soil samples directly adjacent to where the carcass was placed were acquired during the first sampling period for two of the seven sites. Soil samples for the remaining sites were taken 5 m from the carcass during the second sampling period. DNA was extracted from all carcass and soil samples using the PowerSoil DNA Isolation Kit (MO BIO Laboratories, Carlsbad, CA, USA) according to the manufacturer’s instructions and stored at -20°C.

### Bacterial 16S rRNA gene sequencing

The samples were submitted to the Michigan State University genomics core facility for bacterial 16S rRNA gene amplicon sequencing. The V4 region of the 16S rRNA gene (defined by primers 515F/806R) was amplified with dual-indexed Illumina fusion primers as described by Kozich et al. (2013). Amplicon concentrations were normalized and pooled using an Invitrogen SequalPrep DNA Normalization Plate. After library QC and quantitation, the pool was loaded on an Illumina MiSeq v2 flow cell and sequenced using a standard 500 cycle reagent kit. Base calling was performed by Illumina Real Time Analysis (RTA) software v1.18.54. Output of RTA was demultiplexed and converted to fastq files using Illumina Bcl2fastq v1.8.4. Paired-end sequences were filtered and merged with USEARCH 8 (Edgar 2010), and additional quality filtering was conducted with the mothur software platform (Schloss et al. 2009)) to remove any sequences with ambiguous bases and more than 8 homopolymers. Chimeras were removed with mothur’s implementation of UCHIME (Edgar et al. 2011). The sequences were pre-clustered with the mothur command pre.cluster (diffs =1), which reduced the number of unique sequences from 1,136,609 to 784,953. This pre-clustering step removes rare sequences most likely created by sequencing errors (Schloss and Westcott 2011).

### Bacterial diversity analyses

The unique, pre-clustered sequences were considered to be the operational taxonomic units (OTUs) for this study and formed the basis of all alpha and beta diversity analyses, as in our previous study (Dangerfield, Nadkarni, and Brazelton 2017). Sequence reads were not rarefied for alpha diversity and evenness calculations because there was no correlation between diversity indices and sequencing depth for this study. Taxonomic classification of all sequences was performed with mothur using the SILVA reference alignment (SSURefv123) and taxonomy outline (Pruesse, Peplies, and Glöckner 2012). Taxonomic counts generated by mothur and edgeR were visualized using the R package phyloseq 1.20.0 (McMurdie and Holmes 2013).

### Statistical analyses

Alpha diversity and evenness were calculated with the Shannon, invsimpson, and simpsoneven calculators provided in the mothur package (Schloss et al. 2009). Differences between alpha diversities and evenness were tested for significance using the Dunnett-Tukey-Kramer test, which accounts for multiple comparisons among samples with unequal sizes and variances (Lau 2013). Beta diversity was measured using the Morista-Horn biodiversity index, as implemented in mothur. This index was chosen because it reflects differences in the abundances of shared OTUs without being skewed by unequal numbers of sequences among samples. Differences between community compositions were tested for significance using AMOVA (analysis of molecular variance) as implemented in mothur (Pruesse et al. 2012). Morisita-Horn community dissimilarity among samples was visualized using a nonmetric multidimensional scaling (NMDS) plot. This plot was generated using the ordinate and plot ordination commands in phyloseq (McMurdie and Holmes 2013). The ggplot2 function stat_ellipse was added to draw 95% confidence level ellipses (assuming t-distribution) in the NMDS plot (Wickham 2016). Environmental variables (temperature, scavenger counts, and scavenging duration) were fitted to the community composition ordination with the envfit function in the vegan package (Oksanen et al. 2019). Differences in the relative abundance of OTUs between stages was measured using the R package edgeR (Robinson, McCarthy, and Smyth 2009) as recommended by McMurdie and Holmes (2014). The differential abundance of an OTU (as measured in units of log2 fold change by edgeR) was considered to be statistically significant if it passed a false discovery rate threshold of 0.05. Taxa were determined characteristic to that specific stage if they were found differentially abundant in one stage compared to all other stages and soil. These taxa are referred to as “characteristic taxa” for the purposes of this paper. To investigate potential environmental contamination of carcass samples, OTUs with at least 20 sequence counts among all samples were assigned to either fresh carcass or soil using the sink-source Bayesian approach of SourceTracker2 v2.0.1 (Knights et al. 2011) with rarefying to 66,001 sequences for sinks and 16,828 sequences for sources. Similar results were obtained without rarefying sequence counts. The one carcass sample that was determined to be contaminated by soil via SourceTracker2 was excluded from alpha and beta diversity analyses.

### Data Availability

All sequence data are publicly available at the NCBI Sequence Read Archive under BioProject PRJNA525153.

## Results

### Carcass decomposition

Sampling periods were categorized into stages of decomposition based on physical interpretation of the carcasses (Fig 1) as determined from camera trap photographs taken of each carcass and as described by Payne (1965). Day 1 was determined to be “fresh”, as the carcasses were kept frozen promptly after death. The carcasses entered the “bloat” stage in Day 4, as evidenced by the body cavity becoming distended by gases emitted during microbial decomposition. The later sampling periods (Day 12, 18, and 26) were all categorized as the “active decay” stage, because the large decrease is carcass size and the presence of skin tissue on the carcass.

Vertebrate scavengers that fed at the carcasses included American Badger (*Taxida taxus*), Common Raven (*Corvus corax*), Coyote (*Canis latrans*), Kit Fox (*Vulpes macrotis*), Turkey Vulture, and White-tailed antelope squirrel *(Ammospermophilus leucurus*). Turkey vultures were the most frequent scavenger to feed at the carcasses, and the majority of vertebrate scavenging occurred between Day 4 and Day 12 of the study (Table 2).

**Table 1:** AMOVA analysis of significant differences in microbial community compositions. * = significant

| Description | Comparison | F<sub>s</sub> | P-value |
| --- | --- | --- | --- |
| Overall | All 6 sample groups | 9.30736 | <0.001* |
|  | Day 1 vs. Day 4 | 5.88847 | 0.001* |
|  | Day 4 vs. Day 12 | 7.56111 | <0.001* |
|  | Day 12 vs. Day 18 | 0.406973 | 0.738 |
|  | Day 18 vs. Day 26 | -0.22461 | 0.978 |
|  | Soil vs. Day 1 | 11.173 | 0.001* |
|  | Soil vs. Day 4 | 7.47528 | <0.001* |
|  | Soil vs. Day 12 | 14.5593 | <0.001* |
|  | Soil vs. Day 18 | 16.8612 | 0.002* |
|  | Soil vs. Day 26 | 16.8429 | <0.001* |

**Table 2:** Summary of scavenging activity per species per site.

| Site | Species | Individuals | Scavenging Duration (Min) |
| --- | --- | --- | --- |
| Site 1 | Coyote | 1 | 3 |
|  | Turkey Vulture | 1 | 9 |
|  | **Total** | **2** | **12** |
| Site 2 | American badger | 1 | 1 |
|  | **Total** | **1** | **1** |
| Site 3 | Common Raven | 1 | 4 |
|  | Coyote | 1 | 1 |
|  | White-tailed Antelope Squirrel | 6 | 22 |
|  | **Total** | **8** | **27** |
| Site 4 | Common Raven | 3 | 15 |
|  | Kit Fox | 3 | 7 |
|  | Turkey Vulture | 9 | 172 |
|  | **Total** | **15** | **194** |
| Site 5 | Coyote | 2 | 6 |
|  | White-tailed Antelope Squirrel | 1 | 1 |
|  | **Total** | **3** | **7** |
| Site 6 | White-tailed Antelope Squirrel | 1 | 3 |
|  | **Total** | **1** | **3** |
| Site 7 | Common Raven | 16 | 101 |
|  | Coyote | 1 | 1 |
|  | Turkey Vulture | 4 | 85 |
|  | **Total** | **21** | **187** |

### Impact of scavenging on bacterial communities

To identify the impact that scavenging had on bacterial community composition, we compared Morisita-Horn dissimilarities between high and low scavenging sites directly after the peak of scavenging. Sites 4 and 7 were considered to be the “high scavenging” sites as they experienced 88% of the total scavenging duration observed during the study (Table 2). Differences in bacterial community composition between low scavenging and high scavenging sites were not significant, however. Additionally, fitting of scavenging parameters (individuals per week and scavenging duration per week) to the NMDS ordination (Figure 2) yielded no significant correlations.

**Figure 2:**
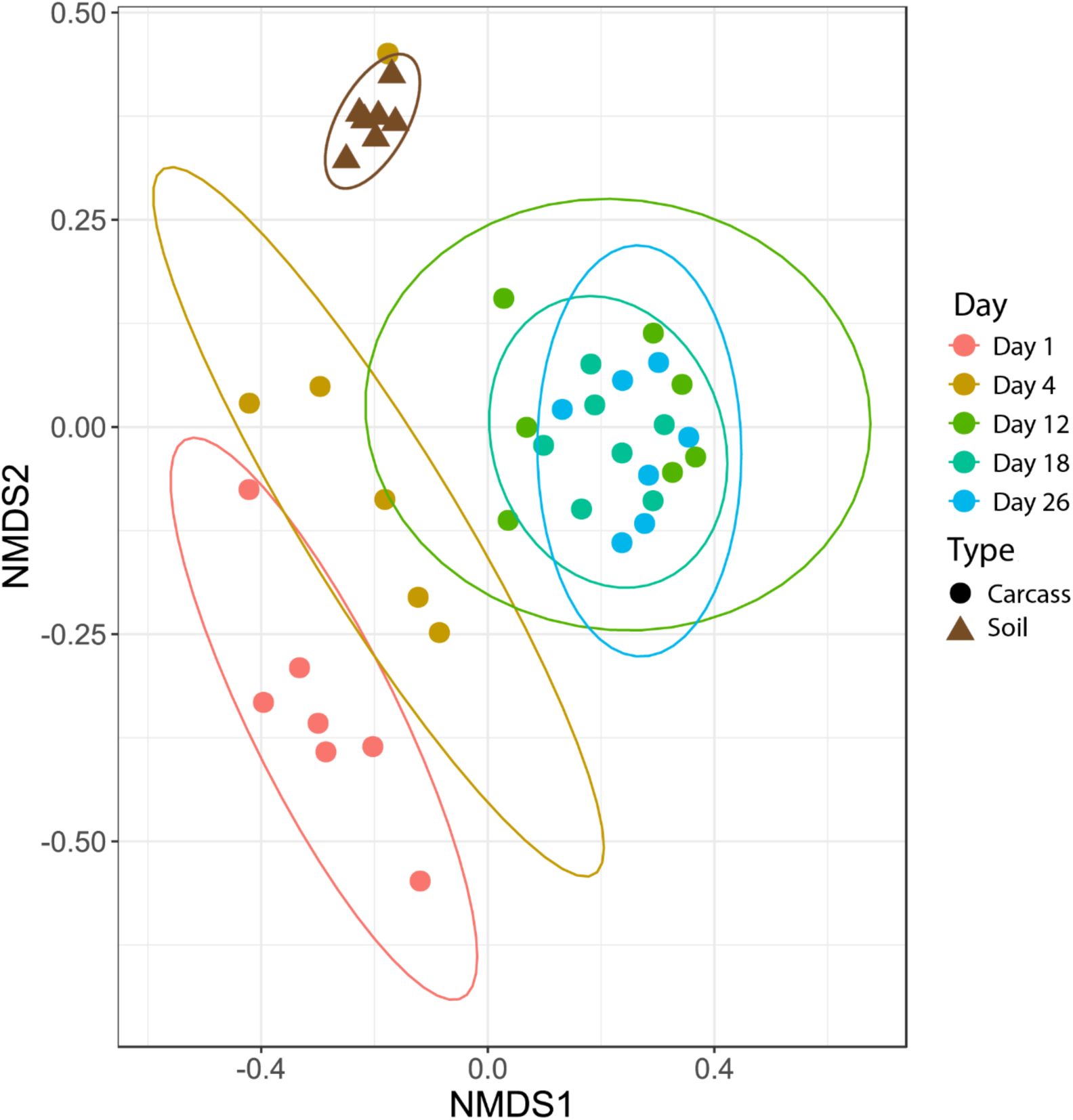
Nonmetric multidimensional scaling (NMDS) plot showing bacteria community shifts associated with the stages of decomposition. The ellipses indicate where 95% of samples within a sample period are expected to occur on the plot.

To identify individual operational taxonomic units (OTUs) that scavengers may have introduced to the carcasses, we contrasted the relative abundances of OTUs in high scavenging sites to low scavenging sites directly after the major scavenging events using edgeR (Robinson et al. 2009). This comparison yielded 39 OTUs that were differentially abundant in high scavenging sites in comparison to low scavenging sites.

We also examined the relative abundance patterns of OTUs classified as genera reported by previous studies to be associated with macroinvertebrates (Dharne et al. 2008; Gupta et al. 2014, 2011; Lee et al. 2014; Shukla et al. 2017; Tóth et al. 2008) All macroinvertebrate-associated genera reach peak relative abundances during the later sampling periods except for Providencia and Myroides (Fig. 3).

**Figure 3:**
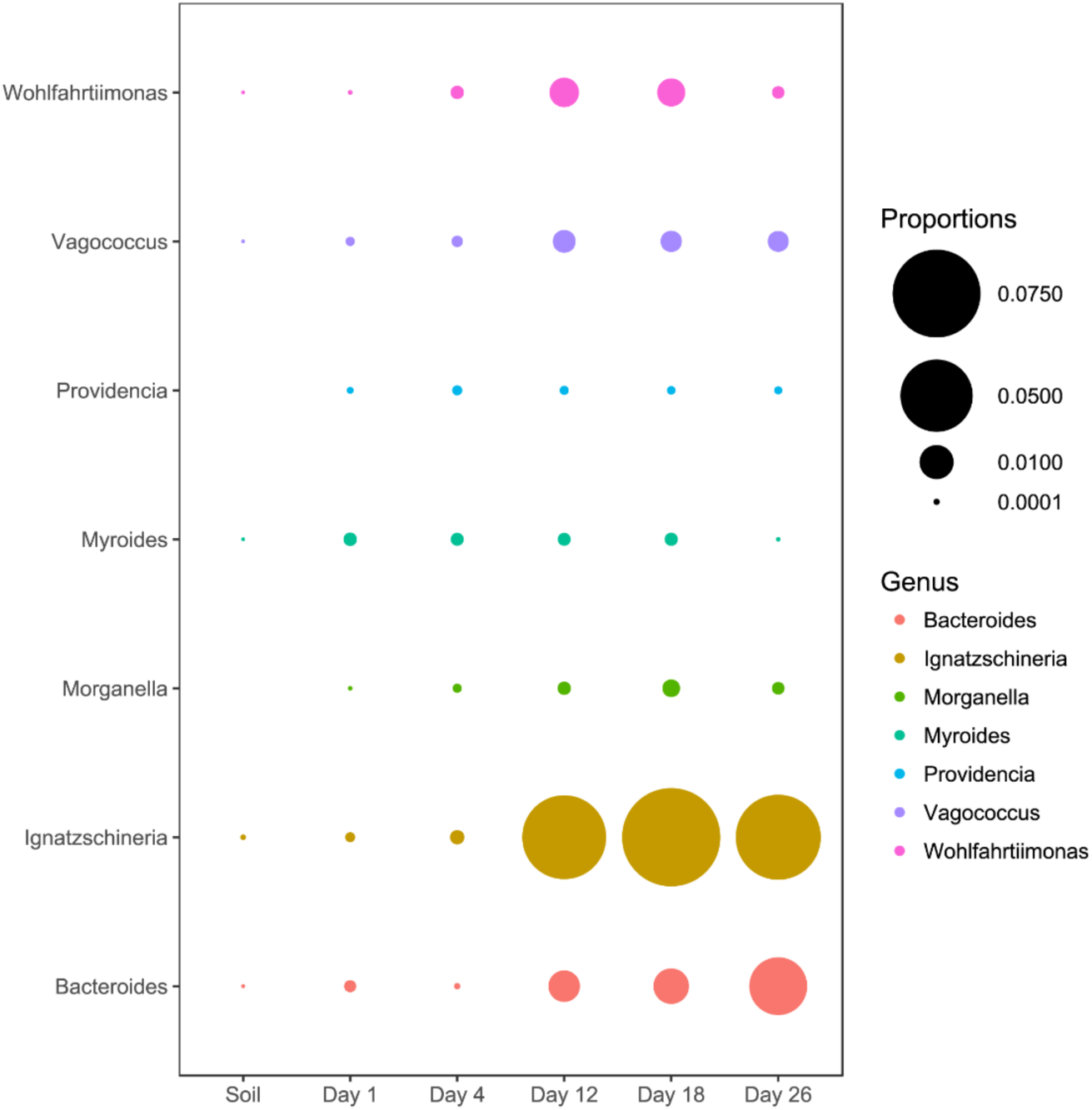
Abundance of genera associated with macroinvertebrates. All genera except Myroides and Providencia exhibited abundance increases in the latter sampling periods.

### Bacterial community changes over time

Microbial community differences are visualized in Fig. 2, where each data point represents the overall bacterial community composition of one sample and the distance between points represents the dissimilarity between samples. There are two primary shifts in community composition: one from Day 1 to Day 4 and a second from Day 4 to later sampling periods (Fig 2). The 95% confidence ellipses show consistent separation between these sampling periods. The only exception to this pattern was a single sample from Day 4 that clusters with soil samples in Figure 2. We examined the bacterial community composition of this outlier sample in more detail with SourceTracker2 (Knights et al. 2011), which revealed that 83% of OTUs in the outlier sample could be confidently assigned to soil. Therefore, we concluded that this sample had been contaminated with soil during sampling and/or handling, and we excluded this sample from all downstream analyses.

Figure 2 also shows that the bacterial community composition of all carcasses at later sampling dates (Days 12, 18, 26) are highly similar and not significantly different from each other. The patterns visualized in the NMDS ordination were tested with an AMOVA that confirmed significant differences between the Day 1, Day 4, and later sampling periods (Table 1). These three clusters of bacterial community composition (Day 1, Day 4, and Days 12-18-26) correspond to the three stages of decomposition identified by physical interpretation of the carcasses (Day 1 = “fresh”, Day 4 = “bloat”, and Days 12-18-16 = “active decay”).

Proteobacteria was the most common phylum in the fresh stage, accounting for 48% of microbial community composition, decreasing to 11% for both the bloat and active decay stages (Figure 4). Conversely, Firmicutes abundance increased as decomposition progressed. Firmicutes increased from 31% in the fresh stage to 72% and 84% of the microbial community in bloat and active decay, respectively. Moraxellaceae represented 30% of the total abundance of the fresh stage (Fig 4), whereas Moraxellaceae only accounted for 2% and 0.8% of the total abundance in bloat stage and active decay, respectively. By contrast, Clostridia was dominant in the bloat stage, accounting for 70% of the total abundance, and accounting for only 3% of total abundance in the fresh stage (Supplementary Krona files).

**Figure 4:**
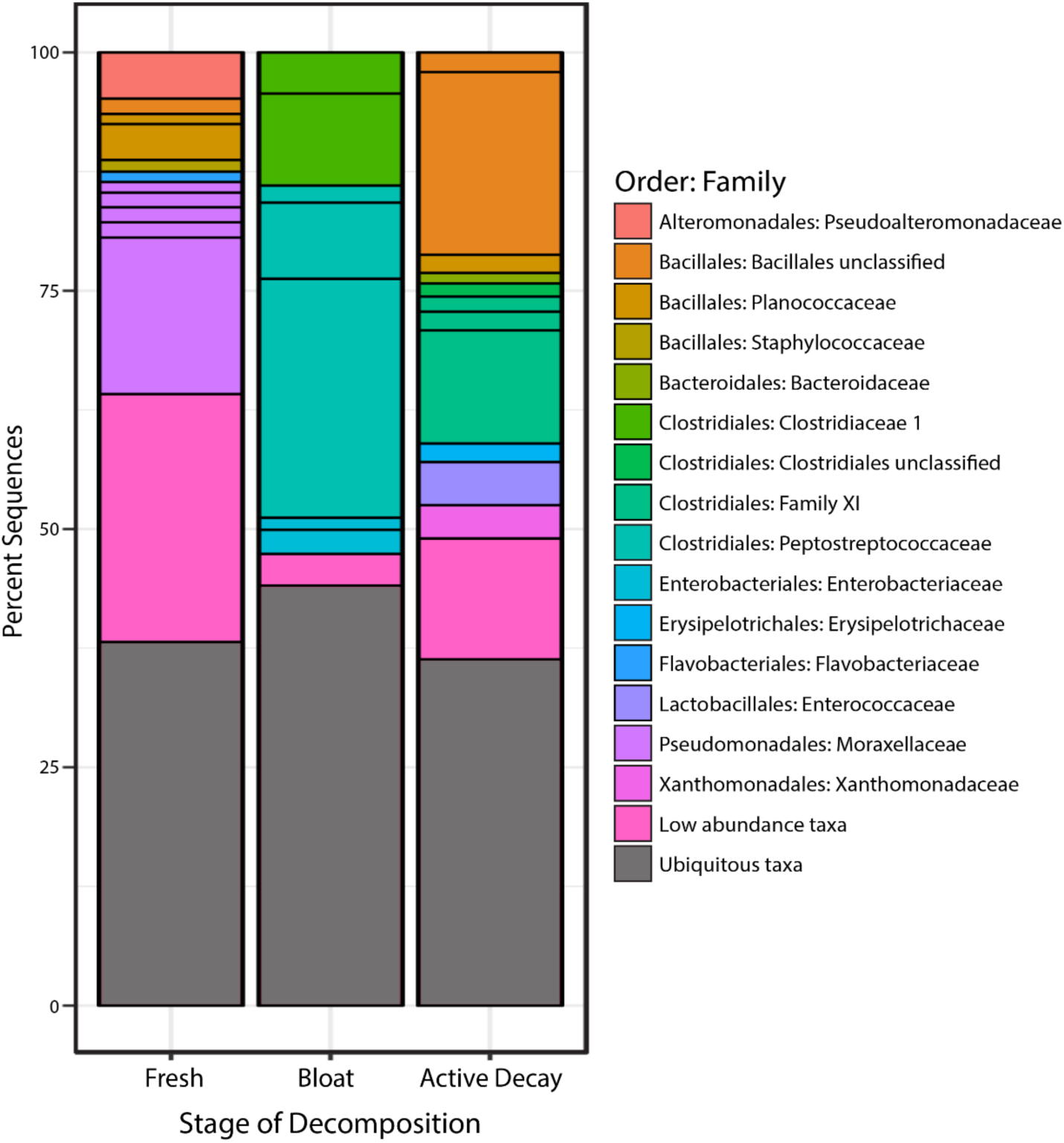
Bacterial communities of each stage of decomposition. Taxa that are present throughout all stages or in soil is represented by “Ubiquitous taxa”. The remain taxa are uniquely abundant within stage of decomposition. Uniquely abundant taxa that had <1% abundance was grouped into “Low abundance taxa”.

### Taxa characteristic to each stage of decomposition

The taxonomic classifications (at order and family level) of all OTUs identified as characteristic to each stage of decomposition (as defined by differential abundance analysis described in Methods) are shown in Figure 4. Each bar in Fig. 4 represents the total community composition of each stage, in which “Ubiquitous taxa” represents taxa that are equally abundant across two or more groups and the remaining taxa in each bar represent OTUs that are characteristic to that stage. There were 323 OTUs that were characteristic to the fresh stage, and these taxa accounted for ∼62% of the total abundance in that stage. These fresh stage taxa are dominated by Moraxellaceae, Flavobacteriaceae, Pseudoalteromonadaceae, Planococcaceae, Staphylococcaceae, and an unclassified Bacillales family. The bloat stage contained fewer characteristic taxa (106 OTUs) than the fresh stage, and these taxa accounted for ∼56% of the total abundance in that stage. The characteristic bloat taxa mainly consisted of Clostridia OTUs. In particular, five Clostridia OTUs accounted for ∼49% of the bloat stage’s total abundance. The active decay stage contained 230 characteristic OTUs that accounted for ∼64 percent of the total abundance. Active decay still contained OTUs from Clostridia, but these OTUs are different than those Clostridia OTUs in the bloat stage and account for less of the total abundance. Active decay stage contained other prominent characteristic taxa from Flavobacteriaceae, Enterococcaceae, Xanthomonadales and an unclassified Bacillales family. All characteristic OTUs, their proportional abundances, and taxonomical classifications are reported in the Supplementary Materials.

### Alpha diversity

Figure 5 reports the OTU Shannon diversity index of each stage of decomposition. Alpha diversity is highest during the fresh stage and decreases during the bloat stage. Diversity increases once the carcass enters the active decay stage. The shifts in Shannon diversity between fresh and bloat stages and between bloat and active decay stages passed a Dunnett-Tukey-Kramer significance test. Similar patterns were also evident with the Inverse Simpson and Evenness (from Simpson) indices.

**Figure 5:**
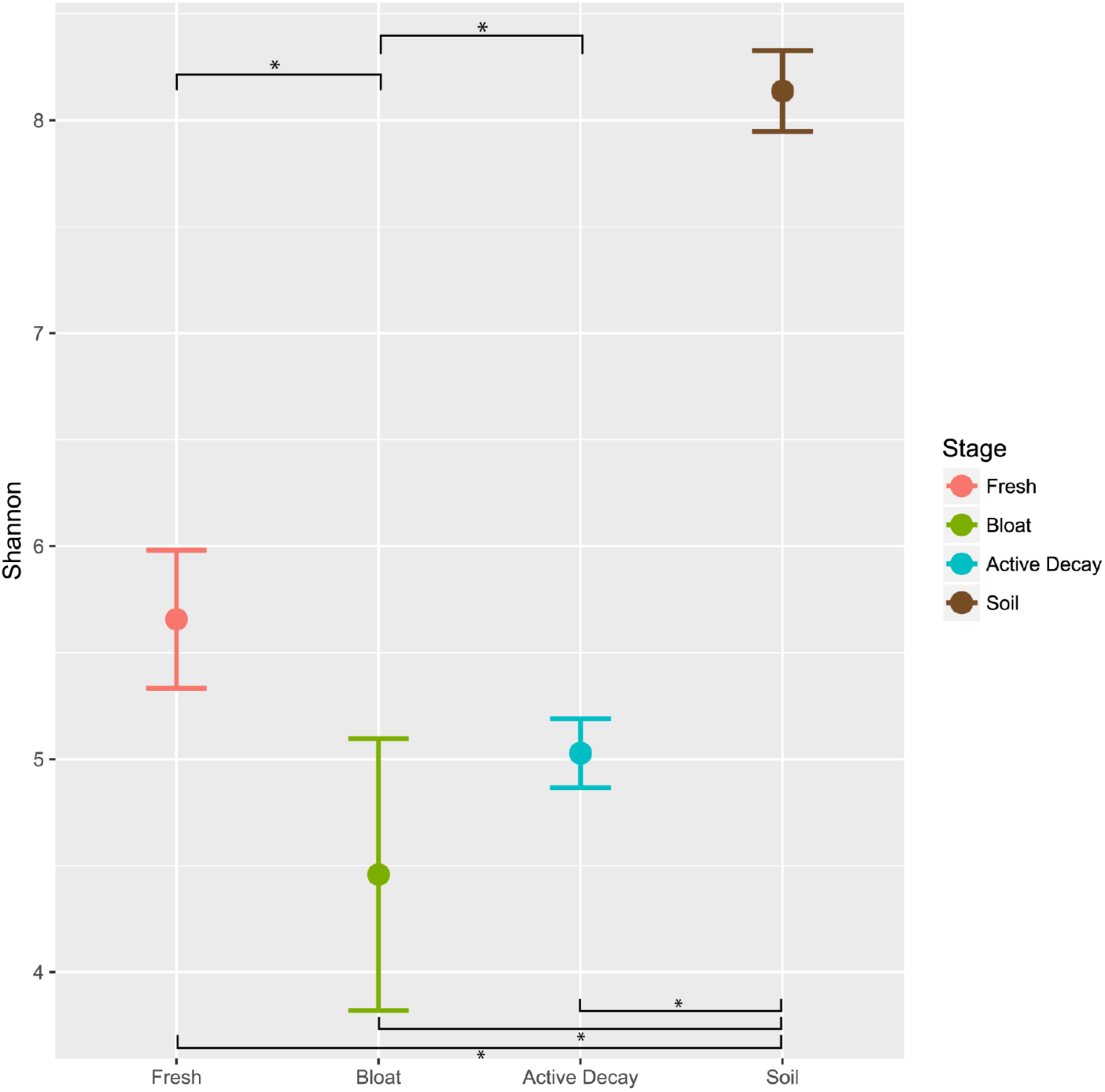
Average Shannon alpha diversity and standard error between decomposition stages and soil. Significance of each comparison was conducted by the Dunnett-Tukey-Kramer test. All comparisons were significantly different from one another except fresh vs active decay. The red dot indicates an outlier (the sample that was contaminated by soil). * = significant (p<0.05)

## Discussion

### No correlation between vertebrate scavenging and bacterial community composition

Carrion is a valuable resource for which many organisms compete, including vertebrate scavengers, macroinvertebrates, and microbes. While there have been some studies documenting the association of bacteria with macroinvertebrates on carrion (Pechal et al. 2013; Rozen et al. 2008; Shukla et al. 2017), this is the first study, to our knowledge, to investigate the impact of vertebrate scavengers on the bacterial community of carrion. Our study site, in an arid region of Utah was well-suited for this experiment because of its unusually sparse vertebrate scavenging activity (Frehner et al. in prep). Studying the succession of bacterial communities in the absence of vertebrate impacts would have been difficult or impossible in a typical environment with vigorous vertebrate scavenging.

Surprisingly, our results indicate no perceptible impact of scavenging activity on the bacterial community composition of carrion. While we did detect 39 OTUs that were more abundant in sites with high scavenging rates, these OTUs were typically present in only a single sample; none of them were consistently present in both high-scavenging sites. Overall, the bacterial communities of carcasses that experienced vigorous vertebrate scavenging (e.g. 15 individuals and 3.2 *hours* of activity) were highly similar to the bacterial communities of carcasses at the same stage of decomposition but with almost no vertebrate scavenging (e.g. 1 individual and 1 *minute* of activity). Additionally, we did not detect any bacterial taxa that were consistently more abundant after peaks in scavenging activity. Because scavenging was so rare at these study sites, it would have been necessary to investigate many more carcasses in order to detect a consistent shift in the bacterial community in response to scavenger activity. Furthermore, the impact of scavenging may be rapid and transient, so future studies should consider sampling at much more frequent time points.

Although we saw no evidence of bacterial taxa consistently associated with vertebrate scavenging, we did find taxa that have been previously associated with turkey vultures (Roggenbuck et al. 2014). However, the abundances of these taxa were not higher in sites that experienced more scavenging by turkey vultures. Furthermore, Roggenbuck et al. (2014) speculated that these taxa are most likely derived not from the turkey vultures, but from the carrion they consume.

### Bacterial taxa associated with macroinvertebrates

Several genera of microbes that have been previously found to be associated with scavenging macroinvertebrates were also present in our study (Dharne et al. 2008; Gupta et al. 2014, 2011; Lee et al. 2014; Shukla et al. 2017; Tóth et al. 2008). With the exception of Providencia and Myriodes, most of these genera exhibited increased abundance when the carcasses entered into active decay. This increase in macroinvertebrate-associated taxa is consistent with the increases of macroinvertebrate activity that occur during the active decay stage(Finley et al. 2015; Matuszewski et al. 2010; Payne 1965).

### Stages of decomposition have consistent bacterial communities

During this study, the carcasses exhibited three stages of decomposition (fresh, bloat, and active decay), and each stage was characterized by a unique community of bacteria. Although the weather, geography, and vertebrate scavenging activity of our study was notably different than in previous studies, our observed patterns of bacterial community succession are remarkably similar to those previously reported for carrion (Hyde et al. 2013; Pascual et al. 2017; Pechal et al. 2014, 2013)Pechal et al. (2013) reported that Proteobacteria was dominant in the early stages of decomposition and declined as decomposition progressed and that Firmicutes abundances progressively increased in the later stages of decomposition. Similarly, 8 of the 14 families that exhibited distinct temporal patterns during the decomposition process in Pascual et al. (2017) exhibited similar patterns in our study (Fig 4).

Our results are also consistent with those of previous studies that have shown a rapid disappearance of aerobic microbial communities during the bloat stage (Finley et al. 2015; Goff 2009; Hyde et al. 2013; Pascual et al. 2017). Abundance patterns of OTUs from Moraxellaceae, all of which are known to be aerobic (Pascual et al. 2017), exemplify this pattern: they represent 30.3% of total abundance in the fresh stage, 2.4% in the bloat stage (when anaerobic Clostridia and Enterobacteriaceae are dominant), and 0.78% in the active decay stage. Furthermore, the dominance of anaerobic bacteria during the bloat stage is exemplified with Clostridia comprising 70% of the microbial community in the bloat stage (Supplementary material bloat krona file). This dominance of bacteria community composition during the bloat stage may partially explain why we observed the bloat stage having the lowest alpha diversity values, whereas Pascual et al. (2017) showed highest alpha diversity during the bloat stage (described as the putrefaction stage).

Most other studies observe more rapid progression through the stages of decomposition (Pascual et al. 2017; Pechal et al. 2014, 2013). It is possible that carcasses during the last sampling period of our study were in “advanced decay”, but there was no discernible change in tissue composition of the carcasses compared to the previous two sampling periods. The delay in decomposition is most likely the result of climatic factors. Previous studies investigating carrion microbial communities were conducted in areas with much higher humidity, whereas this study experienced arid conditions and high temperatures during a period with no precipitation. Active decay has been shown to be limited or hindered by hot and dry climates similar to the conditions present in this study, resulting in partial mummification (Galloway et al. 1989). During this process, carcasses develop a mummified shell over the skeleton as the skin desiccates while macroinvertebrate activity continues underneath, in the body cavity. This partial mummification may explain the prolonged intact composition of the carcass in the latter sampling periods (Fig 1) associated with an increased abundance of macroinvertebrate-associated bacterial taxa (Fig 3).

### Conclusion

In this study, we investigated microbial succession associated with decomposition of cow carcasses that experienced notably little vertebrate scavenging activity. Our results demonstrate that the few intense vertebrate scavenging events at some carcasses did not affect the bacterial community of the carcass. Instead, the bacterial community composition of all carcasses consistently reflected the stage of decomposition, regardless of vertebrate scavenging activity. Our results are remarkably similar to those of other studies conducted in wetter, milder conditions with greater vertebrate scavenging activity (Hyde et al. 2013; Pascual et al. 2017; Pechal et al. 2013, 2014), suggesting that bacterial community succession on carrion follows consistent patterns that are largely unaffected by many external factors.

Although our study did not detect any consistent effects from vertebrate scavengers on whole bacterial communities of carrion, it remains possible that vertebrates could be important vectors of bacteria between carcasses. Such effects are likely to be taxa-specific and may only be detectable under controlled experimental conditions. Future studies could investigate this possibility by sampling carcasses at greater temporal and spatial resolution, by conducting tracing experiments with known vertebrate and microbial taxa, and by characterizing bacterial communities of the skin, mouths, and digestive tracts of vertebrate scavengers.

